# Menger_Curvature : a MDAKit implementation to decipher the dynamics, curvatures and flexibilities of polymeric backbones at the residue level

**DOI:** 10.1101/2025.04.04.647214

**Authors:** Etienne Reboul, Jules Marien, Chantal Prévost, Antoine Taly, Sophie Sacquin-Mora

**Author notes:** Co-first authors (Etienne Reboul, Jules Marien).

## Abstract

Characterizing the dynamics of the backbone of flexible polymers such as Intrinsically Disordered Regions and Proteins (IDRs and IDPs) has proven to be a significant challenge in molecular dynamics (MD) simulations due to their high conformational variability. The widely-used mobility metric Root-Mean-Squared Fluctuations (RMSF) is powerless to provide information for highly flexible systems, as defining a relevant reference structure is often not possible. We previously introduced a new flexibility metric to remedy this gap : the Local Flexibilities (LFs), derived (alongside the Local Curvatures (LCs)) from the Proteic Menger Curvatures (PMCs). Here we present a numba accelerated implementation for any polymer of the calculation of Menger Curvatures as a MDAKit from the widely-used MDAnalysis package. We perform a benchmark with the RMSF and another flexibility metric derived from Proteic Blocks (PBs), the Equivalent Number of PBs (Neq), and show that the PMCs are an order of magnitude faster to compute on a modern CPU chip. We applied all 3 flexibility metrics to a *β*III-tubulin monomer as an example, as tubulins are known to possess the entire range of proteic elements, from *α*-helix and *β*-sheets to a flexible loop and a disordered C-terminal tail (CTT). RMSF, LFs and Neq all succeed in identifying the flexible loops and the CTT, although the RMSF requires a system-specific alignment to do so. Finally, we expose different applications of PMCs, LCs and LFs ranging from mechanism characterization to NMR T2 predictions. We believe that Menger curvatures will prove to be a valuable metric to study protein dynamics and polymers in general. The MDAKit package Menger_Curvature is readily available at https://github.com/EtienneReboul/menger_curvature

## Introduction

New difficulties regarding the analysis of molecular dynamics (MD) simulations have emerged as MD becomes more and more involved in deciphering the behavior of highly flexible polymers such as Intrinsically Disordered Regions and Proteins (IDRs and IDPs). Characterization of their conformational ensemble (CoE) generally relies on global observables such as the radius of gyration. However, this global characterization cannot provide information on the local behavior of the backbone which participates in the IDP or IDR compactness and dynamics. Complex behaviors resulting from transient interactions and strong exposure to the solvent render powerless commonly used metrics such as the Root-Mean-Squared Fluctuations (RMSF). Past attempts to probe or to compare conformational ensembles have proven successful in extending the bioinformatic toolbox available for IDPs and IDRs, but often at the cost of computational speed and interpretability of the metrics [1], [2], [3]. A successful attempt for a simpler flexibility metric at the residue level was already made using Protein Blocks (PBs) by Barnoud et al with their Equivalent Number of PBs (Neq) [4]. By statistically quantifying how many PBs are accessed by an amino acid along a simulation, the Neq can provide a value between 1 and 16 corresponding to low and high flexibility respectively. We recently introduced another metric, the Local Flexibilities (LFs), to successfully characterize the loss in flexibility induced by counterion-mediated bridges in multiphosphorylated peptides [5]. LFs are based on Proteic Menger Curvatures (PMCs), a metric of the backbone curvature at the residue level, and PMCs can also be used to obtain Local Curvatures (LCs) as the average curvature of a residue. To increase performance, applicability, and ease-of-use of the PMCs, we propose a new implementation in the form of a MDAKit named Menger_Curvature, based on the widely-used MDAnalysis ecosystem. We first formally introduce the definition of a Menger curvature, before describing the package implementation. We conclude on a short application to a simulation of a tubulin monomer, compare the results of the LFs to the Neq and the RMSF, and provide a short exposition of successful applications of the metrics to meaningful biological systems.

## Methods

### 1. Definitions of the metrics

#### a. RMSF

RMSF stands for Root-Mean-Squared Fluctuations and is defined by comparing all conformations to a reference, usually the average structure of the molecule or the experimentally resolved structure. The metric is applied to a particle *i* with position *r*_*i*_ with respect to the corresponding particle *j* with position *r*_*j*_ in the reference structure through the following equation :

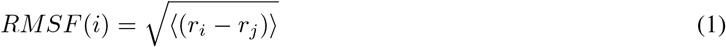

where averaging is performed on all available conformations. In the case of an average structure, *r*_*j*_ is defined as *r*_*i*_ . RMSF usually requires the alignment of the conformations on the reference, which can be challenging if the molecule does not possess stable topological motifs such as secondary structures or bonded segments. Low values of RMSF are usually in the range of 1-2 Å, but its value is technically only bounded by the size of the molecule; the RMSF of an IDP backbone can therefore reach values of several nanometers. The RMSF is a measure of mobility with respect to a reference. In this paper, the reference is arbitrarily defined as the average structure over a trajectory, as is a common practice in the field. The authors would like to warn readers to carefully consider that choice when using this metrics, since an average structure does not always have a biochemical significance, especially in the context of very flexible molecules.

#### b. PBs and Neq

Classification of backbone conformations into Structural Alphabets (SA) can help decipher static and dynamic properties of proteins [6]. A successful example of SA is the Protein Blocks (PB) by De Brevern et al [7]. This alphabet consists of 16 proteic backbone blocks spanning 5 amino acids and labeled with letters “A” to “P”. Dihedral angles are used to define each of them and so, similarly to a Ramachandran plot, each type is usually related to a specific secondary structure or lack thereof. This is discussed in more detail in [4]. Barnoud et al developed a Python-based tool called PBxplore that allows systematic calculations of the PBs and the calculation of a metric called Equivalent Number of PBs (Neq) defined as :

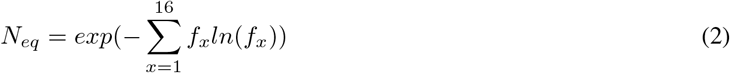

where *f*_*x*_ is the probability of PB *x*. The Neq is bounded by 1 and 16, with values below 2 indicating a very rigid structure, while values above 5 are typically observed for disordered regions and proteins. The Neq is a measure of backbone flexibility [4].

#### c. PMCs, LCs and LFs

Proteic Menger Curvatures (PMCs), Local Curvatures (LCs) and Local Flexibilities (LFs) have previously been introduced by authors to characterize counterion-mediated bridges in multiphosphorylated peptides. These bridges induce hairpins in the proteic backbone, which is therefore less flexible and more curved [5]. The LFs are a measure of backbone flexibility. A visual schematic is available in Figure 1a.

**Figure 1:**
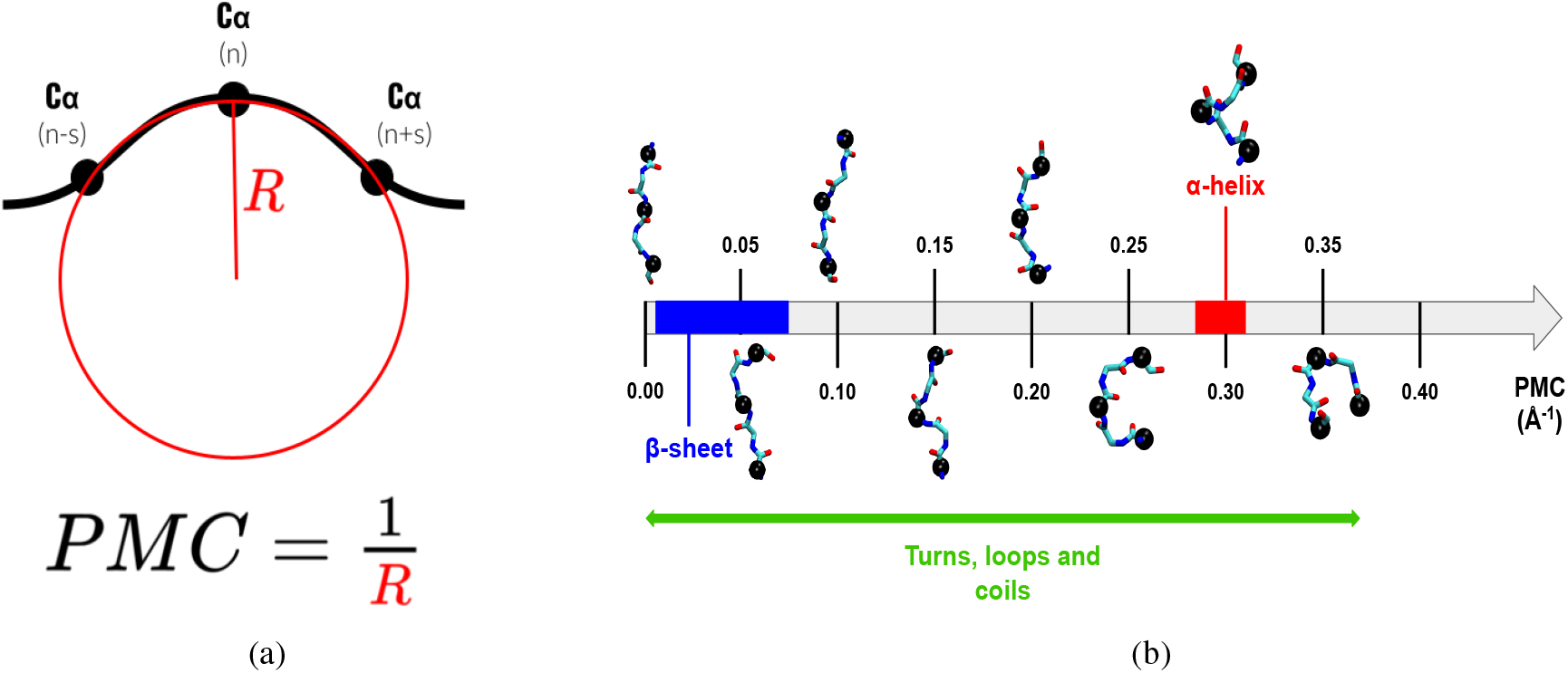
(a) Schematic representation of the Proteic Menger Curvature (PMC) definition. The backbone of the polymer is in black, the *C*_*α*_s are represented by spheres. (b) Range of PMC values and their associated structural elements. Backbone representations are extracted from the single chain tubulin simulation. Backbone is represented in licorice, *C*_*α*_s involved in the PMC calculations are in black Van de Waals. Ranges for *β*-sheets and *α*-helices are based on the ranges of PMC values above 5% in Figure SI-2. These ranges are not meant to be statistically accurate but to provide the reader with a practical estimate of the values that can be expected.x

Consider a triangle. The circumcircle of this triangle (i.e. the circle passing through each of its summits) has a radius *R*, and its inverse is defined as the Menger curvature, named after mathematician Karl Menger [8]. The curvature of a point in a polymer backbone can therefore be defined by selecting two other points of the backbone and calculating the resulting Menger curvature. We here focus on the context of proteins to demonstrate the use of such a metric.

We define the PMC(*n*) by applying the Menger curvature to the triangle formed by the *C*_*α*_ of residue *n* and those of residues *n* −*s* and *n* + *s* where *s* is defined as the spacing. LC(*n*) is the average value of PMC(*n*) over all conformations, and LF(*n*) is the standard deviation over all conformations. The *n* subscript can be dropped for clarity and PMCs, LCs and LFs designate the contracted plural forms of the names. A Python module based on MDAnalysis [9] was previously developed and can be accessed at this Github repository. The aim of this paper is to refine this code, parallelize it and implement it as a fully developed python package as an MDAKit [10]. The resulting Menger_Curvature MDAKit is available at https://github.com/EtienneReboul/menger_curvature.

The choice of spacing *s* is critical as it must be small enough to capture the curvature and flexibility at the residue level but not so small that it becomes redundant with backbone dihedral angles. We previously used a spacing of 2 to successfully characterize the loss of flexibility in a disordered peptide [5]. The persistence length in IDPs can be below a nanometer, indicating that one should not consider lengths longer than 5 amino acids, as the maximum elongation of the backbone would already be of 19 Å [11] [12]. We therefore recommend a spacing of 2 for proteic backbones, but a spacing of 1 might suffice for nucleic acids as their backbone contains twice as many atoms. Increasing the spacing also has the unfortunate consequence of rendering the Menger curvature undefined for the *s* first and last residues, so *s* should be kept as small as possible to retain information at the N- and C-terminals, which are often flexible residues. A spacing of 2 means that the triangle of *C*_*α*_s spans 5 residues as do the PBs, which comforts us that it is a reasonable choice.

### 2. Software dependencies

This project used the cookiecutter template provided for MDAkit . The core of the Menger Curvature package is built upon `MDAnalysis` (version 2.0.0) analysis class [10]. Calculation for the PMC is performed by `NumPy` [13], while `Numba` (version 0.60.0) enables just-in-time compilation for significant performance improvement [14]. Parallel computation is implemented using Python’s standard `multiprocessing` library. For development and testing purposes, we utilized `pytest` [15] for test execution, `pytest-cov` for coverage reporting, and `pytest-xdist` for parallel testing. Documentation was generated using `Sphinx` with the `MDAnalysis` theme.

## Results

### 1. Package implementation

#### a. Installation

Installing the Menger_Curvature package can be done easily with pip inside a conda environment :

~~~
conda create --name menger_curvature
conda activate menger_curvature
conda install pip
pip install --user menger_curvature
~~~

Alternatively, for some High Performance Computers (HPC) the use of conda environment is frowned upon; it is possible to use a virtualenv as a package manager:

~~~
virtualenv menger_curvature
source menger_curvature/bin/activate
pip install menger_curvature
~~~

Another way is to build from source with conda by running the following commands in the folder where you wish to install the program :

~~~
conda create --name menger_curvature
conda activate menger_curvature
git clone https://github.com/EtienneReboul/menger_curvature
cd menger_curvature
conda env update --name menger_curvature --file devtools/conda-envs/test_env.yaml
conda env update --name menger_curvature --file docs/requirements.yaml
pip install -e .
~~~

All package requirements should be taken care of automatically either way. Menger_Curvature is based on MDAnalysis [9], Numpy [13] and Numba [14] for just-in-time compilation.

#### b. Usage and MDAKit architecture

The Menger_Curvature package contains the class MengerCurvature() which allows the calculation of the Menger curvatures of a single-chain polymer backbone with a spacing *s*. We describe how to run the calculation serially with the provided example below.

~~~
# Import the MDAnalysis module
import MDAnalysis as mda
# Import the MengerCurvature class
from menger.analysis.mengercurvature import MengerCurvature
# Create a Universe object with MDAnalysis
u = mda.Universe(“topology.pdb”,
                 “trajectory.dcd”)
# Create the object menger_analyzer
# which will contain the calculations and run # Don’t forget to include the spacing !
menger_analyser = MengerCurvature(u,
                                 “name CA and chainID A”,
                                 spacing=2)
menger_analyser.run()
~~~

If one wishes to run the calculation in parallel, the .**run_parallel()** function can be called. The number of workers can be defined with the **n_workers** variable in the call to **MengerCurvature**, otherwise the class will automatically set **n_workers** to the number of CPUs available on the machine minus 2.

~~~
[…]
menger_analyser = MengerCurvature(u,
                                 “name CA and chainID A”,
                                 spacing=2,
                                 n_workers=10)
[…]
menger_analyser.run_parallel()
~~~

The analysis will return arrays with the Menger curvatures and the associated LCs and LFs. They can be accessed as follows :

~~~
curvatures = menger_analyser.results.curvature_array
local_curvatures = menger_analyser.results.local_curvatures
local_flexibilities = menger_analyser.results.local_flexibilities
~~~

For simulation of *F* frames of a protein with *N* residues, **curvature_array** will have size (*N* −2*s,F* ), **local_curvatures** and **local_flexibilities** will have size *N* −2*s*. The complete architecture of the resulting **menger_analyser** object is available in Figure SI-1. The Menger_Curvature package also comes with a class named **MengerCurvaturePlotter** to properly display the results. A full example is available as a notebook in the documentation : https://github.com/EtienneReboul/menger_curvature/blob/main/notebooks/jm01-tutorial.ipynb).

#### c. Performances

Performances were tested on a small tubulin trajectory of 1000 frames extracted from [16] (450 *C*_*α*_s, last 1000 frames of DCD_protein_ions_trimere_b3_modele16_without_Tau.dcd in https://zenodo.org/records/14888178) and a long trajectory of the Tau protein from [17] (441 *C*_*α*_s, all-atom trajectory of the Tau wild-type in https://doi.org/10.5281/zenodo.14900596).

We provide a table of performances on a laptop with a 14-core 13th Gen Intel(R) Core(TM) i7-13700H processor in Table 1. We compare the implementation of Menger_Curvature and RMSF in MDAnalysis and the Neq calculation with PBxplore.

**Table 1:** Table comparing the computation time in seconds of the PMCs, Neq and RMSF on a short tubulin trajectory (1000 frames, 450 *C*_*α*_s) and a long Tau protein trajectory (20000 frames, 441 *C*_*α*_s). The alignment time of all conformations on the reference was included in the runtime of RMSF as it is necessary to the calculation. Benchmark performed on a 14-core 13th Gen Intel(R) Core(TM) i7-13700H processor.

|  | PMCs, LCs and LFs (serial) | PMCs, LCs and LFs (parallel) | PBs and Neq | RMSF |
| --- | --- | --- | --- | --- |
| Tubulin | $0.23 \pm 0.00$ | $0.19 \pm 0.01$ | $10.54 \pm 0.17$ | $1.21 \pm 0.02$ |
| Tau protein | $4.54 \pm 0.50$ | $2.84 \pm 0.05$ | $218.48 \pm 1.17$ | $22.96 \pm 0.44$ |

We notice that the metrics derived from the Menger_Curvature MDAKit compute at almost the same speed for the short tubulin trajectory. Indeed, the parallelization process is not useful for very small datasets, and the just-in-time compilation with numba takes longer to set up than to run. We therefore recommend users to use the .**run()** method to run the analysis serially in this case. Overall, the performances of the module compare to those of the RMSF, while the computation of the Neq suffers from the lack of parallelization, taking an order of magnitude longer to compute on a standard trajectory than the RMSF or the PMCs.

### 2. Structural and dynamic interpretations

Menger curvature can theoretically span values from 0 to infinity in the unit of the considered system. We will focus on proteins to provide the reader with a better understanding of the metric. Menger_Curvature returns values in units of Å^−1^. Getting an infinite value is, however, not physically possible as steric clashes would never allow for an infinite curvature of the backbone. On the upper range, the maximum value calculated in the applicative case with tubulin is 0.340 Å^−1^ corresponding to a circumcircle of radius 2.942 Å. The maximum value we encountered in IDPs so far in the aforementioned Tau trajectory is 0.371 Å^−1^. Since a value of 0.4 Å^−1^ would correspond to a circumcircle of diameter 5 Å, we can expect that higher values will never be reached as it would be impossible to fit the centers of 3 *C*_*α*_s so close to each other. From our experience, LCs are usually comprised in the range of 0.10 Å^−1^ to 0.30 Å^−1^ for disordered systems and exist at lower values than 0.10 Å^−1^ in loops of structured proteins. LFs can go as low as 0.005 Å^−1^ for very rigid regions, up to 0.030 Å^−1^ in folded cores, and are usually between 0.04 Å^−1^ to 0.10 Å^−1^ for IDPs ([5]). Additionally, the LF value of the central residue of a polyalanine of 13 residues is 0.07 Å^−1^, which could be taken as a reference for a highly flexible region. We present a range of values for PMCs associated to a visual representation of the backbone for values spaced by 0.05 Å^−1^ to help users to better apprehend the metric, and associate secondary structures with the ranges detected in the tubulin simulation (Figure 1b).

*β*-sheets tend to occupy the lower values, mostly from 0.01 to 0.07 Å^−1^, while *α*-helices display a tighter range in the high PMCs, mostly between 0.28 and 0.32 Å^−1^ (Figures 1b and SI-1). One should indeed expect the rigid network of hydrogen bonds to keep *α*-helices from deforming. Turns, loops, and coils can occupy the entire range of possible PMCs, with turns being the only structural elements more curved than *α*-helices (Figure SI-2).

### 3. Applications

#### 3.1 Analysis of a short trajectory of a tubulin monomer

In order to fully demonstrate the capacities of the PMCs, LCs and LFs to characterize the proteic backbone and its flexibility, we needed a system which would display various orders of structuration. Tubulin is a well-studied protein with a very stable core of around 430 amino-acids containing flexible loops necessary to its assembly and its interdimer interactions such as the M-loop (residues 272-286) which interacts strongly with taxol ([18]), and an Intrinsically Disordered Region (IDR) at its C-Terminal Tail (CTT, residues 427-450) ([19]), making it a perfect system to study proteic flexibility. We plotted the LCs and the LFs of a 100ns trajectory (available as test data within the package) to demonstrate the capabilities of the metrics (Figure 2). LCs allow for a simple visualization of the average curvature of the backbone (Figure 2A). LFs are compared to the RMSF and the Neq (Figure 2B). It is clear that LFs are comparable to the Neq as they both correctly identify the M-loop and the CTT as flexible, and that the RMSF concurs with the results of both metrics. Please note that the RMSF was computed on conformations aligned on residues 1 to 426 of the first frame. Excluding the disordered CTT from the alignment is necessary for the relevance of the metric and is archetypal of the bias which is induced when using alignment-based metrics.

**Figure 2:**
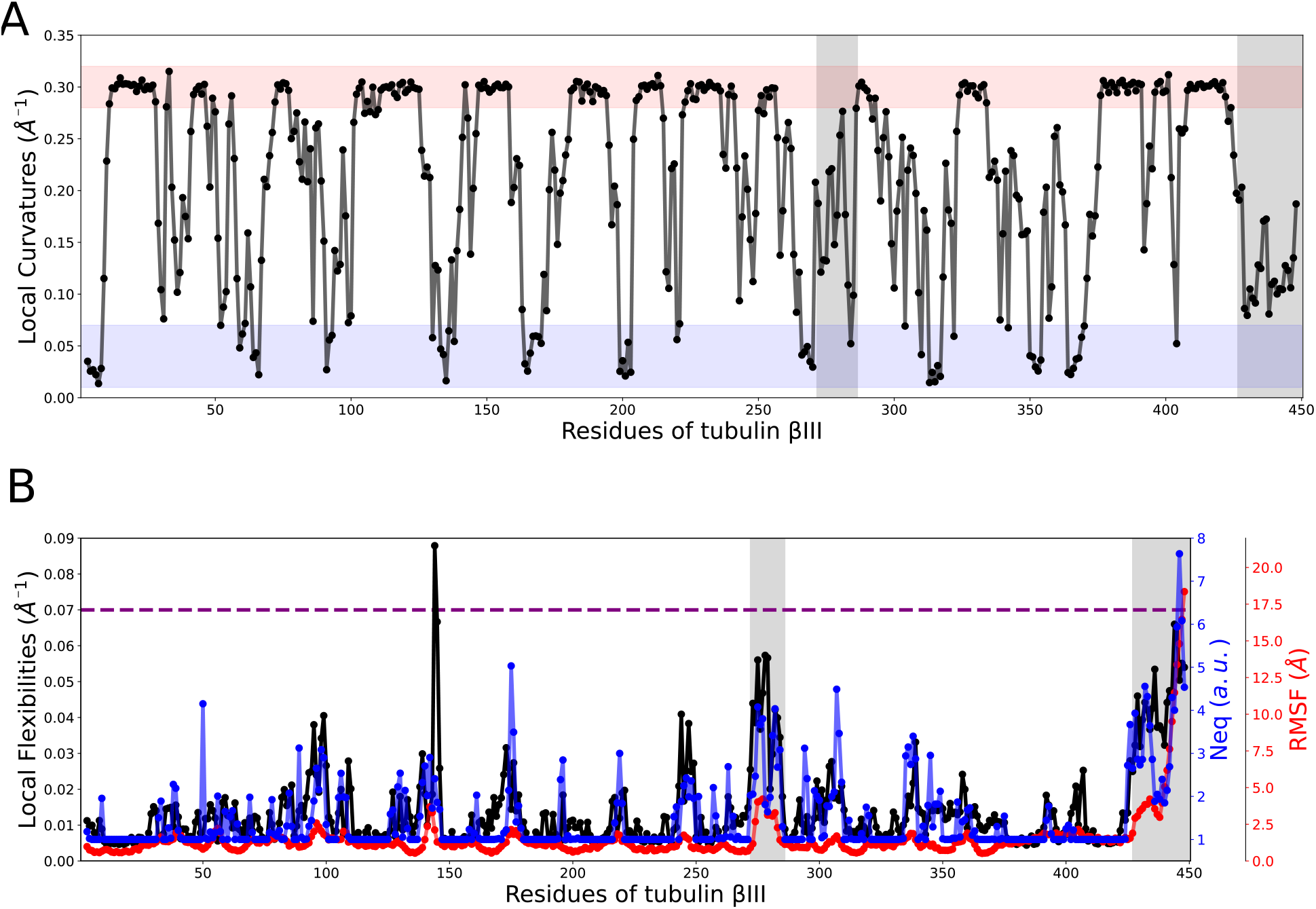
A) Local Curvatures (LCs) of the *β*III-tubulin monomer and B) Comparison of Local Flexibilities (LFs, in black), RMSF with regards to the first structure (in red) and the Neq (in blue). Shaded in grey are the M-loop and the CTT, on the left and on the right respectively. For the LCs, the helical region is shaded in red, the *β*-sheet region in blue. The value of the LFs of a polyalanine 13-mer is traced in purple as a reference (from [5]).

*α*-helices and *β*-sheets can be clearly identified from their very high or very low LCs and low LFs, while the M-loop and the CTT display LFs values 4 to 6 times higher than the rest of the backbone. Residue Gly144 displays a very high LF near 0.09 Å^−1^, which is interesting since it is a part of a turn motif (139-LGGGTG-144) connecting a *β*-sheet to an *α*-helix in the tubulin core, and glycines are known to be very flexible residues due to their absence of side chain. The Neq and the RMSF however do not pick up such a strong difference, highlighting the sensitivity of the LFs compared to the other metrics. Since the goal of this paper is not to be statistically relevant but to provide the reader with insight on the metrics, we will not interpret further this analysis (the complete trajectory of this tubulin is twice as long and several replicas are available in ref [16]).

#### 3.2 Previous successes of the metrics and future perspectives

PMCs, LCs and LFs were first introduced by Marien et al. to characterize the loss of flexibility introduced by a computational artefact in phosphorylated peptides of the Tau protein [5]. LCs and LFs allowed to quantitatively assess that the overbinding of phosphorylations to cations in common all-atom forcefields was causing a bending and a rigidification of the proteic backbone. Later, in a work comparing coarse-grained simulations of the Tau protein, LCs and LFs were shown to concur with EPR data regarding the locality of the pertubation induced by the introduction of phosphorylations at the AT8 epitope of the Tau protein [17]. PMCs were also used to describe along time the “wrapping” phenomenon observed in tubulin C-terminal tails around Tau [16], [20].

But the most striking result obtained with LFs are their correlation to the experimental T2 relaxation time measured in NMR experiments for the Tau protein [21]. Since T2 measurements are often interpreted as a measure of the backbone flexibility, this correlation hints at LFs being more than a mathematical construct, but as a true description of the backbone flexibility. We thus argue that Menger Curvatures and the derived LFs constitute a legitimate progress in the theoretical description of the backbone dynamics of polymers.

Finally, the usage of Menger Curvatures was adapted by Sabei et al to the filament axis of RecA complexes of different lengths [22]. These DNA/protein assemblies, crucial to the homologous recombination process, were shown to exhibit different mechanical properties depending on their size, and the Menger Curvatures and LFs helped characterize this difference and to identify the monomers affected by the change in dynamics.

These example thus highlight that the framework of curvatures applied to molecular assemblies is promising to describe dynamic behaviors for a plethora of systems and scales, and that the Menger_Curvature implementation represents a useful and versatile implementation that can be easily repurposed.

## Discussion and conclusion

A decisive advantage of Menger curvatures is that they are inherent properties of the backbone. As such, they are rototranslationally invariant and the metrics derived from them are not dependent on an alignment like the RMSF. Since each residue but the 2 first and 2 last (if the spacing is set to 2) possesses their own value of curvature independently of whether the structure is part of an ensemble, one can even define a “curvature profile” for each monomer. As a continuous variable, the Menger curvatures could also prove to be useful as a feature in various machine learning applications. We imagine several developments for the Menger curvatures in general, as they could easily be extended to the nucleic acid backbone or to any chain-like system. Possible biopolymers of interest could be RNAs or glycosylated proteins, since the definition of the metric can easily be extended to other atoms or even to branched backbones. It is therefore our opinion that Menger curvatures (and the PMCs in the context of proteins) could become a powerful tool to describe the complex dynamics of polymers, in particular in the context of proteic disorder and folding.

## Data availability

The MDAKit Menger_Curvature is available on Github at : https://github.com/EtienneReboul/menger_curvature

## Acknowledgments

Authors thank Pierre Poulain for fruitful discussions on the Python implementation and the notion of flexibility, Lucas Rouaud for his advice in code development, and Sujith Sritharan and Laetitia Kantin for beta-testing of the MDAKit installation.

## TOC graphic

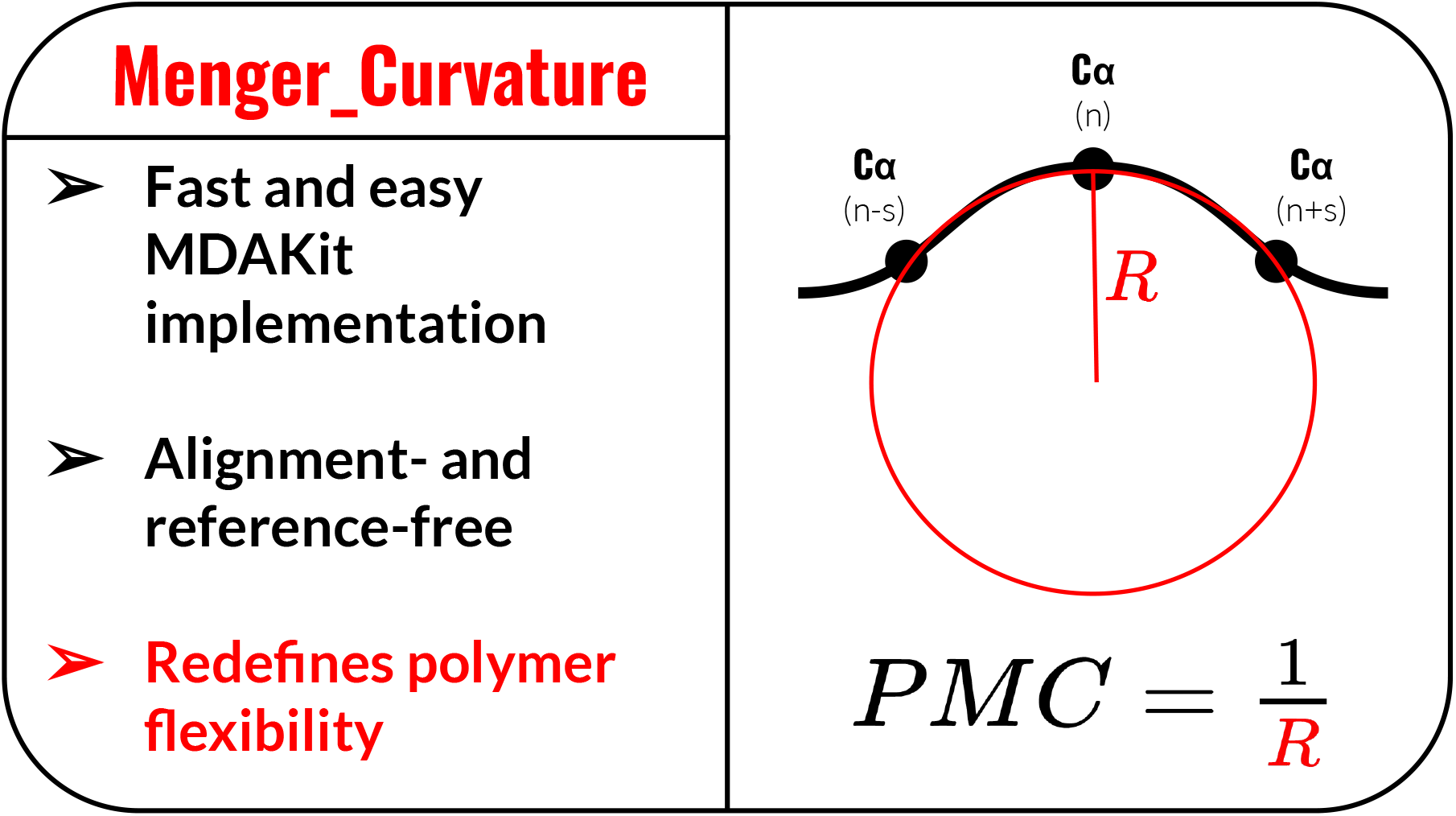

