## Supplementary_Material for "Menger_Curvature : a MDAKit implementation to decipher the dynamics, curvatures and flexibilities of polymeric backbones at the residue level"

September 16, 2026

*Co-first authors
†Corresponding authors

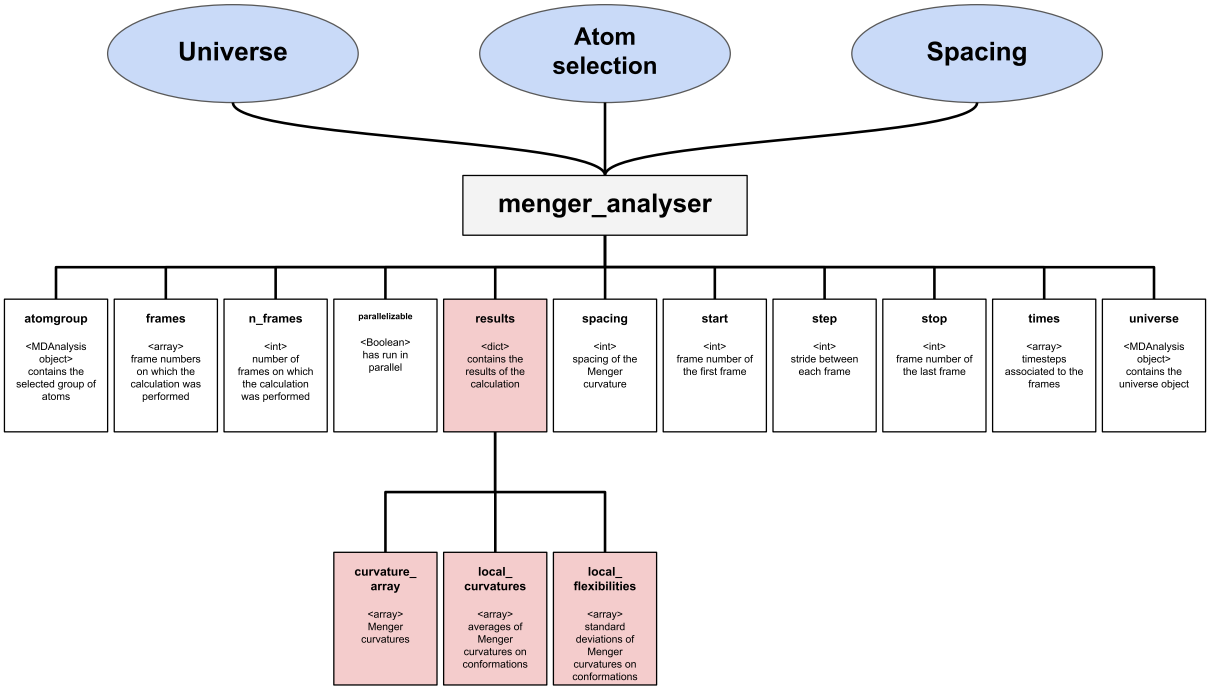


Figure 1: Architecture of the **menger_analyser** object obtained from the **Menger_Curvature** class. Attribute names are in bold, their type is between chevrons followed by a brief description of their content. Required arguments are in blue, results are in red.

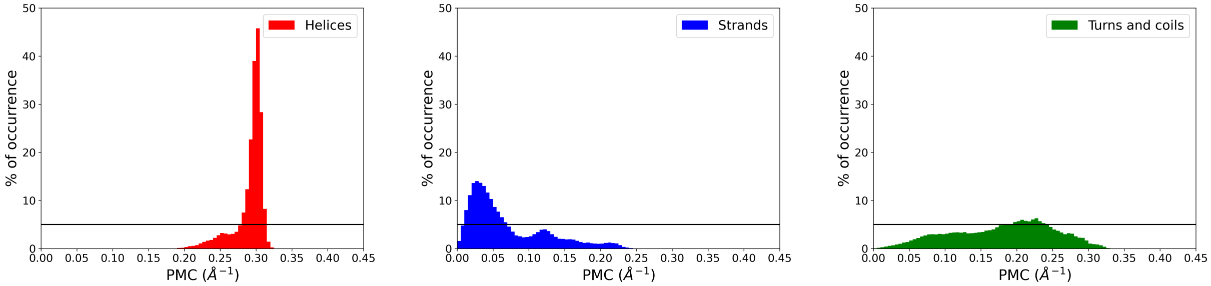


Figure 2: Histogram of probability of PMC values calculated for A) helices, B) strands and C) Turns and coils. The black line signals the 5% threshold. Associated secondary structures were calculated on the short tubulin trajectory with the DSSP algorithm [1] implemented in MDTraj [2].
